# The closed form of Mad2 is bound to Mad1 and Cdc20 at unattached kinetochores

**DOI:** 10.1101/305763

**Authors:** Gang Zhang, Jakob Nilsson

## Abstract

The spindle assembly checkpoint (SAC) ensures accurate chromosome segregation by delaying anaphase onset in response to unattached kinetochores. Anaphase is delayed by the generation of the mitotic checkpoint complex (MCC) composed of the checkpoint proteins Mad2 and BubR1/Bub3 bound to the protein Cdc20. Current models assume that MCC production is catalyzed at unattached kinetochores and that the Mad1/Mad2 complex is instrumental in the conversion of Mad2 from an open form (O-Mad2) to a closed form (C-Mad2) that can bind to Cdc20. Importantly the levels of Mad2 at kinetochores correlate with SAC activity but whether C-Mad2 at kinetochores exclusively represents its complex with Mad1 is not fully established. Here we use a recently established C-Mad2 specific monoclonal antibody to show that Cdc20 and C-Mad2 levels correlate at kinetochores and that depletion of Cdc20 reduces Mad2 but not Mad1 kinetochore levels. Importantly reintroducing wild type Cdc20 but not Cdc20 R132A, a mutant form that cannot bind Mad2, restores Mad2 levels. In agreement with this live cell imaging of fluorescent tagged Mad2 reveals that Cdc20 depletion strongly reduces Mad2 localization to kinetochores. These results support the presence of Mad2-Cdc20 complexes at kinetochores in agreement with current models of the SAC but also argue that Mad2 levels at kinetochores cannot be used as a direct readout of Mad1 levels.

## INTRODUCTION

The spindle assembly checkpoint (SAC) is the major regulator of chromosome segregation throughout eukaryotes and generates a “wait anaphase” signal in response to unattached kinetochores^1,2^. The biochemical nature of this “wait anaphase” signal is the mitotic checkpoint complex (MCC) composed of the checkpoint proteins Mad2 and BubR1/Bub3 (Mad3 in yeast) bound to the protein Cdc20 ^3–5^. Cdc20 is the mitotic coactivator of the anaphase-promoting complex (APC/C) and the MCC blocks the activity of APC/C-Cdc20 hereby preventing anaphase onset^6^. Current models propose that unattached kinetochores act as catalysts for MCC production in agreement with the fact that all checkpoint proteins localize to unattached kinetochores and that laser ablation of unattached kinetochores allows progression into anaphase^7–10^. The rate-limiting step in MCC production is the binding of Mad2 to Cdc20 as this requires a large conformational change in Mad2 from an open form (O-Mad2) to a closed form (C-Mad2)^11–14^. Closed Mad2 binds to short similar sequences in both Cdc20 and Mad1 but the interaction with Mad1 is stable while Mad2 can be removed from Cdc20 via the p31^comet^/TRIP13 pathway^15–20^.

The Mad1/Mad2 complex at kinetochores catalyzes the conformational conversion of Mad2 and binding to Cdc20. Specifically the Mad1/Mad2 complex binds a O-Mad2 molecule through Mad2 heterodimerization and the C-terminal domain of Mad1 catalyze the conversion to C-Mad2 by an unknown mechanism^11,14,21–23^. The proximity of Cdc20 to the Mad1/Mad2 complex mediated by the Bub1 checkpoint protein further facilitates the binding of Mad2 to Cdc20 ^11,14,24–26^. Given the central role of the Mad1/Mad2 complex in catalyzing the rate-limiting step in SAC signaling it is not surprising that the strength of the SAC in cells correlates with Mad2 levels at kinetochores^27,28^. However it is not fully established if C-Mad2 at kinetochores only reflects the levels of Mad1/Mad2 or if kinetochore localized Cdc20 also contributes. This is important to establish, as it will have implications for the interpretation of Mad2 kinetochore signals in terms of molecular composition.

Here we use a monoclonal antibody specific for C-Mad2 and live cell imaging of fluorescently tagged Mad2 to explore the contribution of Cdc20 to Mad2 kinetochore localization.

## RESULTS

### The kinetochore levels of C-Mad2 and Cdc20 correlate

We synchronized HeLa cells using a thymidine block and release protocol combined with nocodazole treatment (Figure 1A). Nocodazole is a spindle poison that activates the SAC by generating unattached kinetochores. Cells were fixed and stained with our recently described C-Mad2 specific monoclonal antibody that stained unattached kinetochores (Figure 1B)^29^. We noted a larger variation in the levels of C-Mad2 compared to Mad1 (Figure 1C). The level of C-Mad2 on kinetochores in the cell with the highest C-Mad2 kinetochore signals is almost 100 times more than the level in the cell with the lowest C-Mad2 kinetochore signals while it is only 4 times for Mad1. We reasoned that a population of C-Mad2 at kinetochores might be in complex with Cdc20 and therefore we co-stained cells for Cdc20 and C-Mad2. Indeed we visually observed a clear correlation between Cdc20 kinetochore levels and C-Mad2 levels (Figure 1B). Quantification of the levels of Cdc20 and C-Mad2 on single kinetochores and normalizing this to CREST levels confirmed a correlation between their levels (Figure 1D).

**Figure 1.**
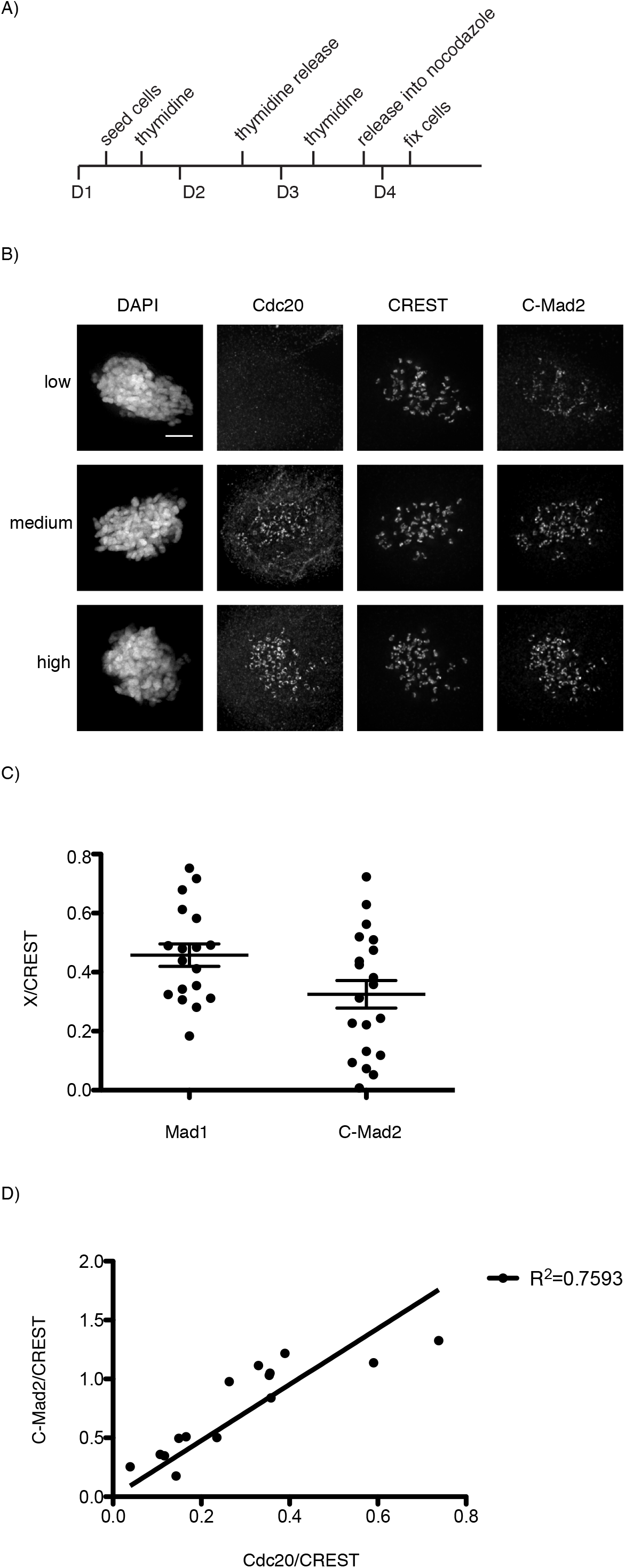
Kinetochore signals of Cdc20 and closed Mad2 are highly correlated. A) Protocol used for B). B) Overnight nocodazole treated cells were fixed and co-stained by C-Mad2 and Cdc20 antibodies together with CREST antibody. Three cells expressing different levels of Cdc20 and C-Mad2 are presented. Bar is 5 μm. C) Nocodazole treated cells were fixed and stained with either Mad1 antibody or C-Mad2 antibody and CREST antibody. 18 cells for Mad1 and 20 cells for C-Mad2 were randomly picked for quantification of kinetochore levels of Mad1 or C-Mad2 and normalized to CREST level. A single dot represents the average kinetochore level from 20 kinetochores in one cell. D) The kinetochore signals of C-Mad2 and Cdc20 are normalized to CREST signals and the correlation is presented. Each dot represents a single cell. 10 pairs of kinetochores in each cell are quantified. Scale bar, 5 μm.

These results support that a fraction of C-Mad2 at kinetochores could be in complex with Cdc20.

### Mutating the Mad2 binding site in Cdc20 lowers C-Mad2 kinetochore levels

We reasoned that the variation in Mad2 levels were the result of variation in the time cells had been arrested in prometaphase. To achieve a more homogenous population of prometaphase cells we arrested cells in G2 using the Cdk1 inhibitor RO3306 (Figure 2A). Cells released from RO3306 into medium with nocodazole enter mitosis shortly after and we fixed and stained cells for C-Mad2 and observed a much more homogenous staining of kinetochores. To analyze the role of Cdc20 in C-Mad2 kinetochore localization we efficiently depleted Cdc20 using RNAi and co-stained cells for Cdc20 and either Mad1 or C-Mad2 (Figure 2B-E). While Mad1 levels were unaffected by Cdc20 depletion we observed an approximately 50% decrease in C-Mad2 levels. To confirm that the reduction in C-Mad2 kinetochore levels upon Cdc20 depletion was due to removal of kinetochore localized Mad2-Cdc20 complex we complemented cells with RNAi resistant Cdc20 constructs. While wild type Cdc20 restored C-Mad2 levels to control levels a point mutant, Cdc20 R132A, unable to bind Mad2 did not (Figure 2F-G). In agreement with previous work showing that BubR1 and Bub1 localize Cdc20 to kinetochores the depletion of Mad2 did not affect Cdc20 kinetochore levels (Figure 2H-I).

**Figure 2.**
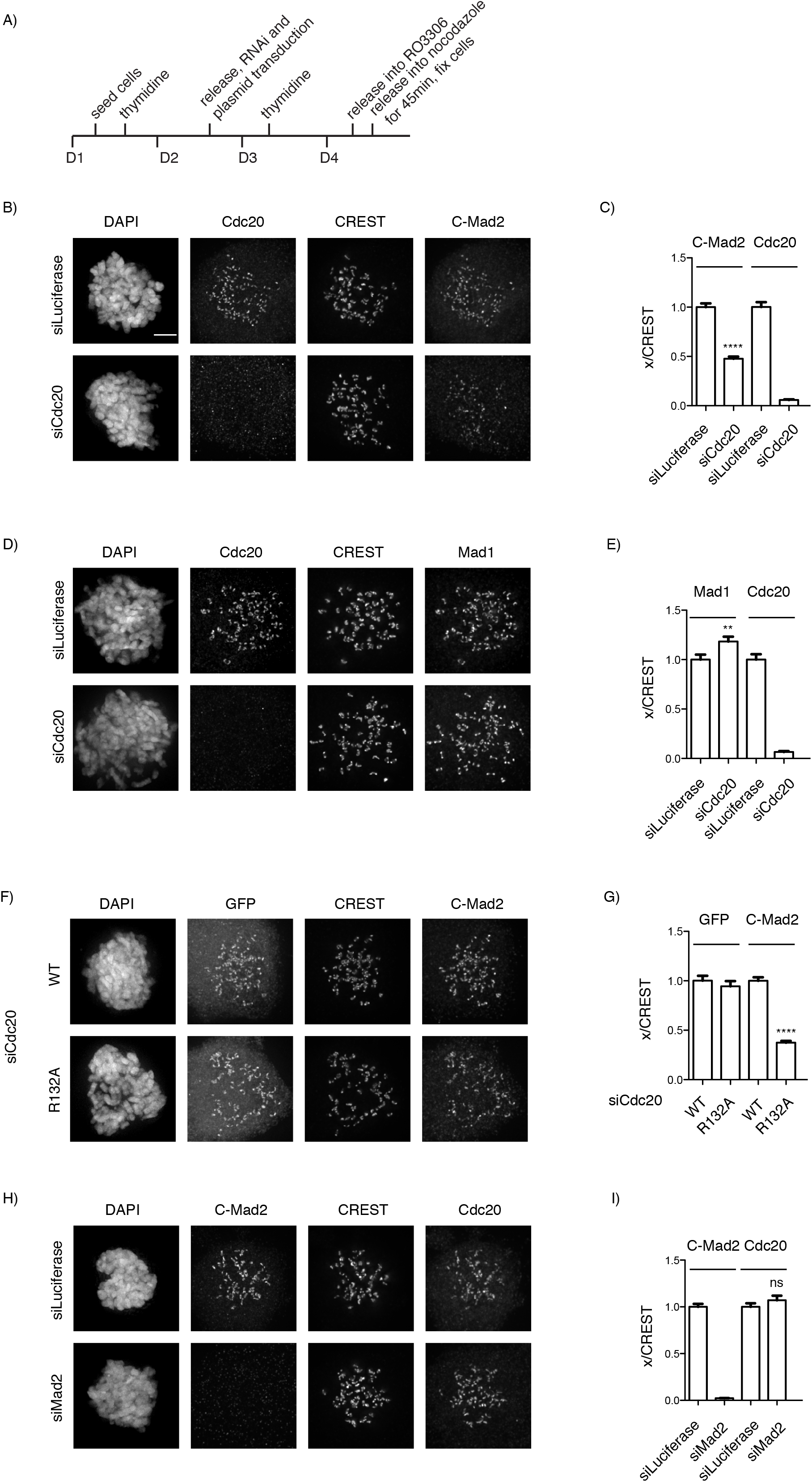
C-Mad2 kinetochore localization depends on Cdc20. A) Optimized protocol for B-I). B) siRNA oligos against luciferase or Cdc20 were transfected into HeLa cells. After nocodazole teatment for 45 min, cells were fixed and costained with Cdc20, C-Mad2 antibodies together with CREST antibody. Bar is 5 μm. C) 100 pairs of kinetochores from 10 cells from A) were quantified and plotted. D) Similar treatment as in B) was performed. Cells were co-stained with Cdc20, Mad1 antibody together with CREST antibody. E) 100 pairs of kinetochores from 10 cells from D) were quantified and plotted. F) siRNA against Cdc20 together with RNAi resistant constructs for either wild type Cdc20 or R132A Cdc20 mutant were co-transfected into HeLa cells. Similar treatment as in B) was performed except GFP antibody instead of Cdc20 antibody was used. G) 100 pairs of kinetochores from 10 cells from F) were quantified and plotted. H) siRNA oligos against luciferase or Mad2 were transfected into HeLa cells. After nocodazole treatment for 45 min, cells were fixed and co-stained with C-Mad2, Cdc20 antibodies together with CREST antibody. To avoid cells exit mitosis due to depletion of Mad2, MG132 was added together with Nocodazole. I) 100 pairs of kinetochores from 10 cells from H) were quantified and plotted. A Mann Whitney test was used for statistical comparison of the different samples. (**** : P ≤ 0.0001, ns: non significant). Scale bar, 5 μm.

To provide further evidence that Mad2 kinetochore levels represents in part its complex with Cdc20 we expressed Mad2-Venus in HeLa cells and filmed cells as they progressed through mitosis. In control depleted cells we observed clear kinetochore localization of Mad2-Venus while Cdc20 depletion strongly inhibited Mad2-Venus localization (At least 30 cells for each condition analyzed showing similar Mad2 localization) (Figure 3).

**Figure 3.**
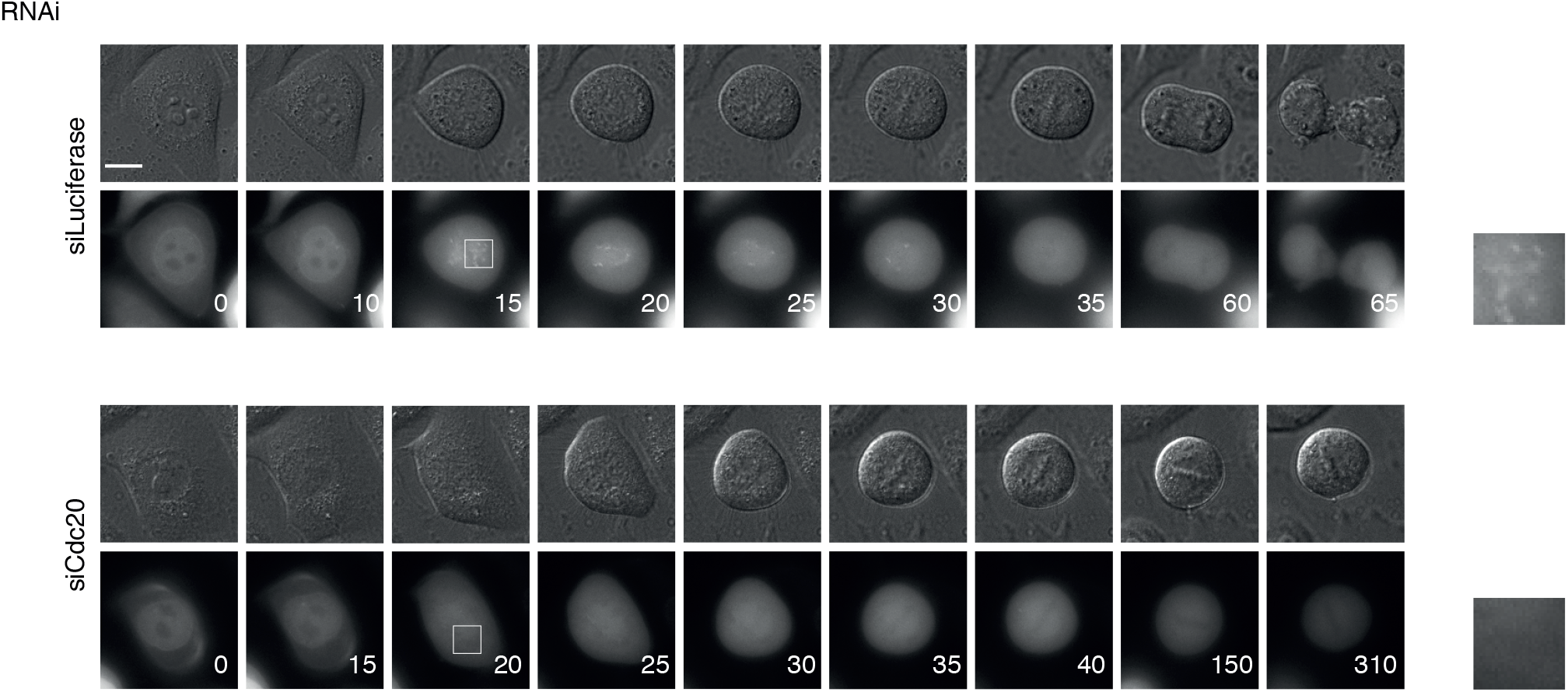
Live cell analysis of Mad2-Venus localization in Cdc20 depleted cells. A Mad2-Venus plasmid was transfected into HeLa cells before RNAi of Cdc20 or luciferase. Live cell imaging was conducted on the cells to monitor the kinetochore levels of Mad2-Venus when cells entered mitosis. Scale bar, 5 μm. At least 30 mitotic cells per condition frpm 2 independent experiments were analyzed.

Combined our data show that C-Mad2 at kinetochores is not only in complex with Mad1 but also Cdc20.

## DISCUSSION

The role of Mad1/Mad2 as a catalyst for Mad2-Cdc20 complex formation is well established and our data support that C-Mad2 kinetochore signals represent both the catalyst and product. This is important to keep in mind when interpreting Mad2 kinetochore levels in molecular terms and our data argue that Mad2 kinetochore levels cannot be used as a direct readout of Mad1 levels. Previous data have also shown that Cdc20 removal reduces the levels of a Mad2 mutant protein that cannot bind p31^comet^ although in these experiments no effects were observed on exogenous myc-tagged wild type Mad2 and off target effects of Cdc20 RNAi were not accounted for^17^. Interestingly a previous analysis with a different C-Mad2 specific antibody revealed that 50% of kinetochore localized C-Mad2 is in complex with p31^comet 30^. Based on the results presented here this could reflect p31^comet^ binding to C-Mad2 in complex with Mad1 or Cdc20.

Currently we do not know what is causing the variation in Mad2-Cdc20 complex levels at kinetochores but we assume that this relates to the time cells have spend in mitosis because we achieve high and homogenous Mad2 levels when cells are captured shortly after release from Cdk1 inhibition. We favor that upon entry into mitosis there is an initial burst of MCC production at kinetochores, reflected by high Mad2-Cdc20 complex levels at kinetochores, but following this a low level of MCC formation is sufficient to maintain the SAC.

## MATRIALS AND METHODS

### Cell culture

HeLa cells were cultivated in DMEM medium (Invitrogen) supplemented with 10% fetal bovine serum and antibiotics. Cells were synchronized by thymidine (2.5 mM) the day before co-transfection with siRNA oligos against Cdc20 or Mad2 (50 nM as final concentration) and corresponding plasmids by Lipofectamine 2000 (Life Technologies). RNAi oligos targeting Cdc20 (5’ CGGAAGACCUGCCGUUACAUU 3’) or luciferase (5’ CGUACGCGGAAUACUUCGA 3’) or Mad2 (5’ GGAAGAGUCGGGACCACAG 3’), were used for RNAi depletions. Thymidine was added again 12 hours later after transfection. 24 hours later, the cells were released from thymidine block into medium with RO3306 (10 nM as final concentration). 8 hours later, cells were released from RO3306 into fresh medium containing nocodazole (200 ng/ml as final concentration). Fixation was performed after 45 min as described above.

### Immunofluorescence and quantification

Cells growing on coverslips were synchronized with a double thymidine block and RO3306 (10 nM) block. After washing with PBS, the cells were treated with medium containing nocodazole (200 ng/ml) for 45 min and fixed with 4% paraformaldehyde in PHEM buffer (60 mM PIPES, 25 mM HEPES, pH 6.9, 10 mM EGTA, 4 mM MgSO_4_) for 20 minutes at room temperature. Fixed cells were extracted with 0.5% Triton X-100 in PHEM buffer for 10 minutes. The antibodies used for cell staining include Cdc20 (Bethyl, A301-180A-1, rabbit, 1:200), CREST (Antibodies Incorporated, 15-234, human, 1:400) and GFP (Abcam, ab290, rabbit, 1:400). Mad1 (Santa Cruz, sc-65494, mouse, 1:200), C-Mad2 (home made, mouse, 1:200) All the fluorescent secondary antibodies are Alexa Fluor Dyes (Invitrogen, 1:1000). Z-stacks 200 nm apart was recorded on a Deltavision Elite microscope (GE Healthcare) using a 100X oil objective followed by deconvolution using Softworx prior to quantification. Protein intensity on kinetochores was quantified by drawing a circle closely along the rod-like CREST staining covering the interested outer kinetochore protein staining on both ends. The intensity values from the peak three continuous stacks were subtracted of the background from neighboring areas and averaged. The combined intensity was normalized against the combined CREST fluorescent intensity.

### Live cell imaging

15 ng per well of pcDNA5/FRT/TO Mad2-Venus plasmid was transfected into HeLa cells grown in 6-well plate 48 hours before live cell imaging. After 24 hours, the transfected cells were treated with thymidine and re-seeded into Ibidi dish (Ibidi). RNAi against Cdc20 was performed in Ibidi dish overnight. Cells were released from thymidine block and medium changed to Leibovitz’s L15 medium (Life technologies) supplemented with 10% FBS before filming. Live cell imaging was performed on a Deltavision Elite system using a 40 x oil objective (GE Healthcare).

## ACKNOWLEDGEMENTS

We thank members of the lab for discussions and support.

